# Proliferative and quiescent human gastric cancer stem-like cells are associated with differential chemoresistance and patient mortality

**DOI:** 10.1101/2020.10.23.351726

**Authors:** Kok Siong Ang, Hong Kai Lee, Marion Chevrier, Michelle Goh, Jingjing Ling, Vivien Koh, Hang Xu, Xiaomeng Zhang, Jia Chi Tan, Nicole Yee Shin Lee, Kelvin K. L. Chong, Sergio Erdal Irac, Ze Ming Lim, Josephine Lum, Alicia Tay, Charles Antoine Dutertre, Carolyn Tang, Calista Wong, Mai Chan Lau, Chun Jye Lim, Samuel Wen Jin Chuah, Sherlly Lim, Joe Poh Sheng Yeong, Valerie Chew, Anis Larbi, Amit Singhal, Msallam Rasha, Yoshiaki Ito, Michael Poidinger, Matthew Chau Hsien Ng, Shanshan Wu Howland, Patrick Tan, Florent Ginhoux, Jimmy So, Wei Peng Yong, Jinmiao Chen

**Author notes:** Corresponding author Jinmiao Chen, 8A Biomedical Grove, Immunos, #03-06 Singapore 138648, Wei Peng Yong, 5 Lower Kent Ridge Road, NUH Medical Centre (NUHMC) Levels 8-10, Singapore 119074. These authors contributed equally (Kok Siong Ang, Hong Kai Lee). These authors contributed equally (Marion Chevrier, Michelle Goh).

## Abstract

**Objective:** Gastric cancer (GC) tumors are highly heterogenous with different subpopulations of epithelial cells. We employed single cell RNA sequencing (scRNA-seq) to dissect the heterogeneity and identified subpopulations of cancer cells with stem-like properties. We further investigated their resistance to oxaliplatin chemotherapy and their contribution to gastric cancer outcome.

**Design:** We performed scRNA-seq on FACS sorted epithelial and immune cells from paired samples of GC tumors and normal adjacent tissues. We identified two epithelial subpopulations (STMN1^+^IQGAP3^+^ and STMN1^+^IQGAP3^−^) with stem-like properties. We characterized and compared them to known healthy gastric stem cell populations. We also cultivated GC derived organoids to study the chemoresistance of similarly marked populations. Lastly, we employed immunohistochemistry (IHC) staining to ascertain the predicted immunosuppressive interactions.

**Results:** The STMN1^+^IQGAP3^+^ subpopulation showed a higher tumor mutation burden, upregulated proliferative pathways and transcriptomically resembled proliferative healthy gastric isthmus stem cells. The STMN1^+^IQGAP3^−^ subpopulation were comparatively quiescent and transcriptomically resembled enteroendocrine cells. Both transcriptomic signatures were associated with worse mortality than other epithelial subpopulations with the quiescent being associated with the poorest patient survival.

GC tissue derived organoids were dominated by STMN1^+^IQGAP3^+^ cells but the STMN1^+^IQGAP3^−^ compartment was more resistant to chemotherapy. We also verified the likely suppression of CD8 T cell cytotoxicity by STMN1^+^IQGAP3^+^ cells through the NECTIN2/TIGIT interaction.

**Conclusions:** Cancer cells with stem-like characteristics are associated with poor survival through chemoresistance and immunosuppression. Reactivating the immune system through checkpoint blockade is an opportunity to eliminate these cells.

**What is already known on this topic:** Multiple gastric stem cell populations have been identified and linked to tumor initiation in rodent-based studies. However, none of them have been conclusively proven in human tumors. Isolating and characterizing tumor cells with stem-like properties will help shed light on their possible origin and possible mitigation strategies.

**What this study adds:** Here we identified two sets of stem-like gastric cancer cells that are associated with poorer patient prognosis. One set is highly proliferative and exhibits oxaliplatin susceptibility. It also engages in immunosuppressive interactions such as NECTIN2/TIGIT. The other set is quiescent and highly resistant to oxaliplatin.

**How this study might affect research, practice or policy:** The transcriptome signatures of the identified stem-like cells can aid in patient prognosis and identify patients who can benefit from checkpoint blockade therapy to reactivate their immune response towards gastric cancer cells.

## Introduction

Gastric cancer (GC) is currently the fifth most common cancer and the third cause of death by cancer globally [1]. Surgical resection followed by chemotherapy are standard of care with immunotherapy, neoadjuvant, and even radiotherapy are being explored to improve outcomes [2]. Without proactive screening, GC is usually discovered at late stages with poor survival prospects. The resistance of these advanced tumors to standard chemotherapy poses additional major challenges [3]. Moreover, checkpoint inhibitor therapies focused around PD-1/PD-L1 interaction has not been a resounding success in treating GC [4,5]. There is still a great need to understand the mechanisms for relapse and therapy resistance in GC patients to guide intervention [5].

Therapy resistance and eventual relapse has been partially attributed to small populations of cancer stem cells (CSCs) in many cancers [6]. CSCs are long-lived tumor cells with self-renewal abilities, and they have been detected in various leukemia and solid tumors [7]. CSCs can seed relapse and metastasis, as a few cancer stem cells are enough to initiate new tumors [8]. Various CSCs have also been reported to suppress CD8 T and NK cell functions [9] and resist apoptosis [10]. For highly proliferative and plastic CSCs, random mutations accumulate and many CSCs have been shown to be chemotherapy resistant [11]. Thus, CSCs are an important target for successful cancer treatment.

CSCs can originate from stem cells gaining cancerous properties or cancer cells gaining stemness [12,13]. CSCs developing from stem cells can arise from random mutations [14] or aberrant stroma signaling and stress from inflammation [15]. Various stem cell populations have been reported at the gastric pits in the antral and corpus regions in the stomach of healthy lab mice. In the antral glands, stem cells are marked by Lgr5 [8,9,15] at the base, and different stem cell populations are marked by Bmi1^+^ [17], Mist1^+^ (or Bhlha15) [18], eR1^+^ [19], Cckbr^+^ (Cck2r) [20], and AXIN2^+^ [21] at the isthmus. In the stomach corpus glands, the pit base harboured different chief cell populations marked by Lgr5 [16], Troy [22], and Mist1 [23] that function as reserve stem cells that are activated by injury. At the corpus glands isthmus, Mist1 [24] and Bmi1 [17] mark two quiescent stem populations, while eR1 [19], Iqgap3 [25], Stmn1, and Mki67 [26] mark an actively proliferating stem cell population responsible for corpus epithelial homeostasis.

Among the myriad of gastric stem cells discovered, some, such as the Lgr5^+^, Mist1^+^, and Iqgap3^+^ populations, had been proposed to be the origin of gastric cancers based on lineage tracing experiments [18,25,27]. However, these studies employed mouse models to study tissue repair/renewal and cancer initiation, and the process can differ in humans. Gastric cancer organoids offer another approach to study CSCs; such organoids have been derived from CSCs in human GC cell lines [28,29] and patient samples [30,31]. These studies take place after *in vitro* expansion, thus out of the context of the tumor environment. Our current understanding of human gastric stem cells is also lacking, with cancer development trajectories presently unclear.

Single cell RNA-sequencing (scRNA-seq) technology is well-suited to study the rare cancer stem cells from human patient samples. Zhang *et al*. applied scRNA-seq on gastric pre-malignant lesions and early intestinal GC tissue, identifying metaplastic stem-like cells (MSCs) overexpressing OLFM4, EPHB2, and SOX9 [32]. These cells were found in intestinal metaplasia lesions and increased in proportion with the progression into early gastric cancer. However, in-depth characterization of these MSCs and links to known healthy gastric stem cells are still lacking, which should shed light on gastric cancer’s origin in humans and treatment options. There is a need to further characterize potentially diverse human GC CSC populations and how they possibly developed from their healthy counterparts. Furthermore, we aim to understand the mechanisms for chemo resistance and immune resistance in GC CSC populations, and single cell technologies enable us to identify the responsible cell populations and interactions [33].

In this work, we used FACS and Smart-seq2 for scRNA-seq analysis of tissues from 15 histologically classified GC patients (i.e., diffuse, intestinal, and mixed) at different GC stages. We characterized the epithelial and immune cells with a multimodal bioinformatics approach and found two stem-like subpopulations (STMN1^+^IQGAP3^+^ and STMN1^+^IQGAP3^-^) that correlated with poor patient prognosis. STMN1^+^IQGAP3^+^ cells bore transcriptional similarity to the proliferative isthmus stem cells [25] and were named proliferative stem-like epithelial cells (pSE). STMN1^+^IQGAP3^-^ cells were similar to enteroendocrine cells at the bottom of gastric glands and were named quiescent stem-like epithelial cells (qSE). In patient-derived GC organoids, STMN1^+^IQGAP3^-^ qSEs were more resistant to oxaliplatin than proliferative STMN1^+^IQGAP3^+^ pSEs, and patients bearing the qSE signature showed the poorest post-chemotherapy survival. We also found that pSE cells expressed various immunosuppressive ligands inhibiting CD8 T and NK functions.

## Methods

### Patient samples

The research activities involved in this study were reviewed and approved by the local institutional ethics board (National Healthcare Group Domain Specific Review Board: B/2016/00059 and B/2005/00440) and complied with local laws and regulations. The study was conducted in accordance with the Declaration of Helsinki and International Conference on Harmonization and Good Clinical Practice guidelines. All tissue samples were obtained with written informed consent from all patients involved. The patients underwent total or subtotal gastrectomy at the National University Hospital of Singapore. They were eligible for study enrolment if they had a histologically or cytologically confirmed diagnosis of primary gastric cancer, were 21 years or older, and had never undergone prior chemotherapy or radiotherapy. Patients with symptomatic or progressive central nervous system metastases, or other uncontrolled medical disorders were ineligible.

Matching tumor-edge and normal tissues were collected prospectively from GC patients who visited the National University Hospital in Singapore between July 2016 and May 2017. All tissue samples were obtained from surgical specimens immediately following removal from patients undergoing primary surgery, with no pre-operative irradiation or chemotherapy. Briefly, 6-8 surgical bites (each approximately 5 mm in size) on each of the fresh gastric tumor and normal-adjacent tissues were collected from the patients during planned surgical resection. The bites were preferably obtained from the periphery of the resected tissue whenever possible to capture regions known to contain higher proportions of immune cells. Gastric adenocarcinoma was confirmed by histological examination and then classified according to Lauren’s criteria. Demographics and clinicopathological profiles of the patients are summarized in Supplemental Table S1. Detailed materials and methods are described in the online supplemental methods.

## Results

### Dissecting gastric cancer heterogeneity at single cell resolution

Paired gastric tissue samples (tumor and normal adjacent) were collected from 15 GC patients, with six patients diagnosed histologically as diffuse, four as intestinal, and five as mixed (Table S1). Using Smart-seq2 in conjunction with index sorting, we performed single-cell full-length transcriptome sequencing and obtained raw data of 3443 cells (Figure 1A). We then employed the Seurat package [34] for downstream analysis. After quality filtering, 2878 cells were retained with a median of 2342 genes detected per cell (Figure 1B). We identified 12 cell clusters (Figure 1C) and manually annotated them by marker expressions (Table S2). We identified fibroblasts, epithelial cells, and different immune cells including natural killer (NK) cells, T cells, B cells, plasma cells, and macrophages among the sequenced cells.

**Figure 1:**
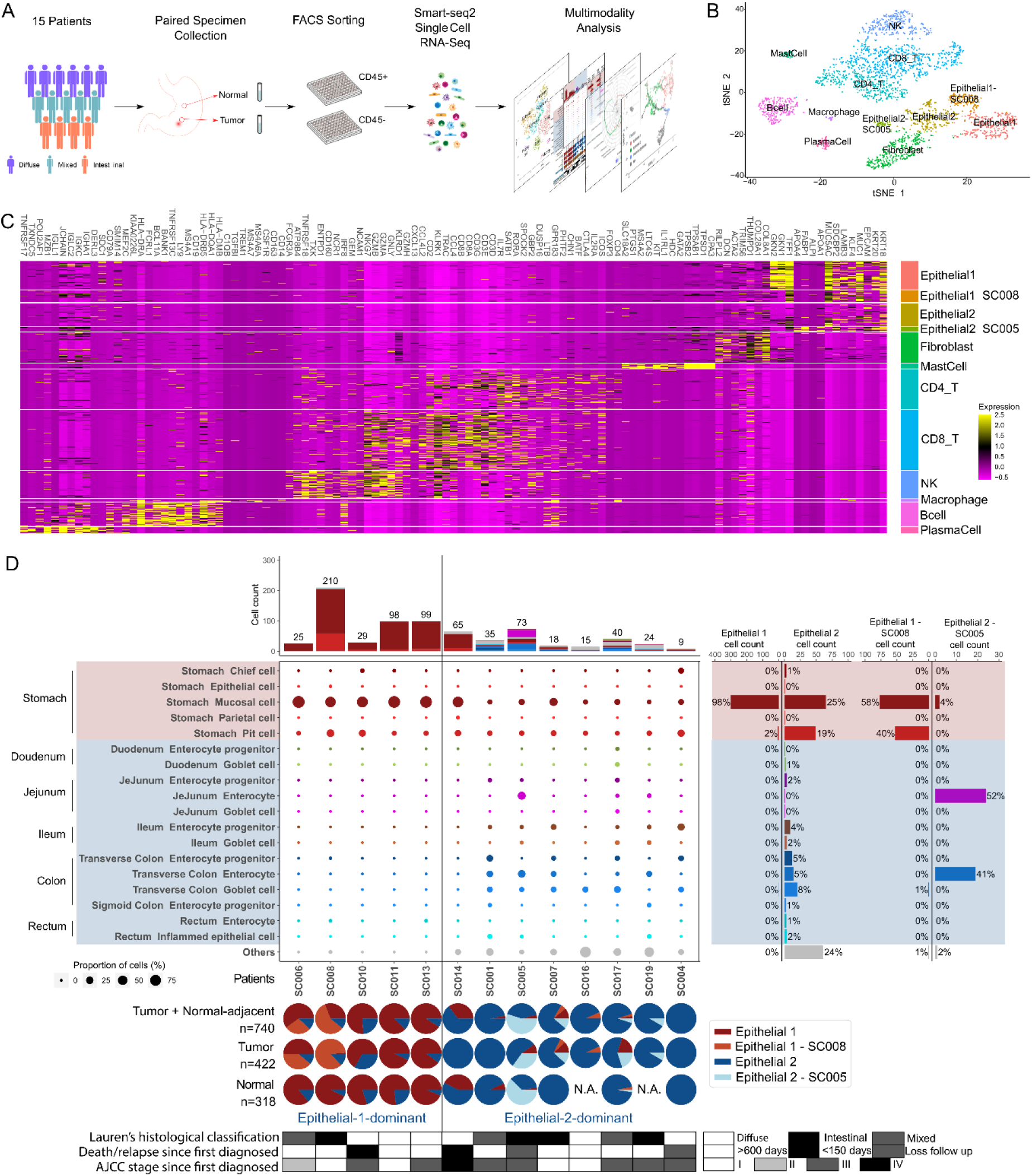
Tumor and normal tissues collected from gastric cancer patients can be classified into Epithelial-1-dominant (Gastric-dominant) and Epithelial-2-dominant (Gastrointestinal-mixed) tissues. A) A schema of overall study workflow. B) Overview of the 2878 cells that were selected (QC-passed) for subsequent analyses in this study (tSNE). C) Differential expression genes for cell-type clusters identified in (B). Each row represents one cell and each column represents a marker gene known for the respective cell-type cluster. D) The scHCL predicted closest matching organs/ cell-types of the epithelial cells obtained from each patient. The middle panel shows the closest matching organ/cell-type of the epithelial cells (row) by patient (column). The size of circle indicates the proportion of epithelial cells for each specific organ/cell-type. The circles are color coded by the closest matching organs and cell types, respectively. The histogram on the top shows the cumulative number of each organ/cell-type-specific cells, grouped by patient. The histogram on the right panel shows the count and proportion of organ/cell-type-specific cells found in the respective clusters identified in (B), including Epithelial 1, Epithelial 2, Epithelial 1 specific to Patient SC008, and Epithelial 2 collected from Patient SC005. The pie chart tracks show proportions of epithelial clusters found in tumor and/or normal tissues collected from each patient. The bottom annotation tracks show (from top to bottom): the Lauren’s histological classification, number of days to death since first diagnosed, and AJCC stage since first diagnosed.

We found two major clusters of epithelial cells, labelled as Epithelial 1 (E1) and Epithelial (E2) [35]. The E1 cells largely originated from patients with diffuse GC and resembled gastric cells by cell type prediction with ScHCL (https://github.com/ggjlab/scHCL), while E2 cells mainly originated from patients with intestinal GC and resembled intestinal cells (Figure 1D) [36]. There are two subclusters dominated by cells from patients SC008 (E1-SC008, 92%) or SC005 (E2-SC005, 87%) (Figure 1C). Patient SC008 was clinically diagnosed as a well-differentiated Stage IA adenocarcinoma while SC005 was diagnosed with rare mucinous adenocarcinoma. SC005 is notable with a unique mix of predicted cell types unlike all other patients; most of the cells were predicted to be jejunum- and colon-like. These characterizations illustrate the utility of single cell sequencing in dissecting the inter and intra tumoral heterogeneity to identify subtypes.

### Identification of stem-like epithelial cells associated with poorer patient survival

Next, we isolated the epithelial cells and re-clustered them for in-depth characterizations. We observed six different epithelial clusters: the previously identified E1, E1–SC008, E2 and E2– SC005 clusters, and two new E2 derived clusters that harbored stem-like properties (Figure 2A, Table S3). Gene regulatory network analysis with SCENIC [37] first revealed four clusters based on their regulon activities (Figure S2A). Most of the E1 and E2 cells fell into the Regulon clusters 0 and 1, while 93% and 48% of the two new clusters were in Regulon cluster 2. In Regulon 2, we observed high activity among the stem cell-related transcription factors, such as CDX1, MYC, and HDAC1 (Figure S2B; Table S4). On the gene expression level, the new clusters had several cancer stem cell markers upregulated, including OLFM4 [32], CD44 [38], PROM1 [39], and ALDH1A1 [40] (Figure S2C).

**Figure 2:**
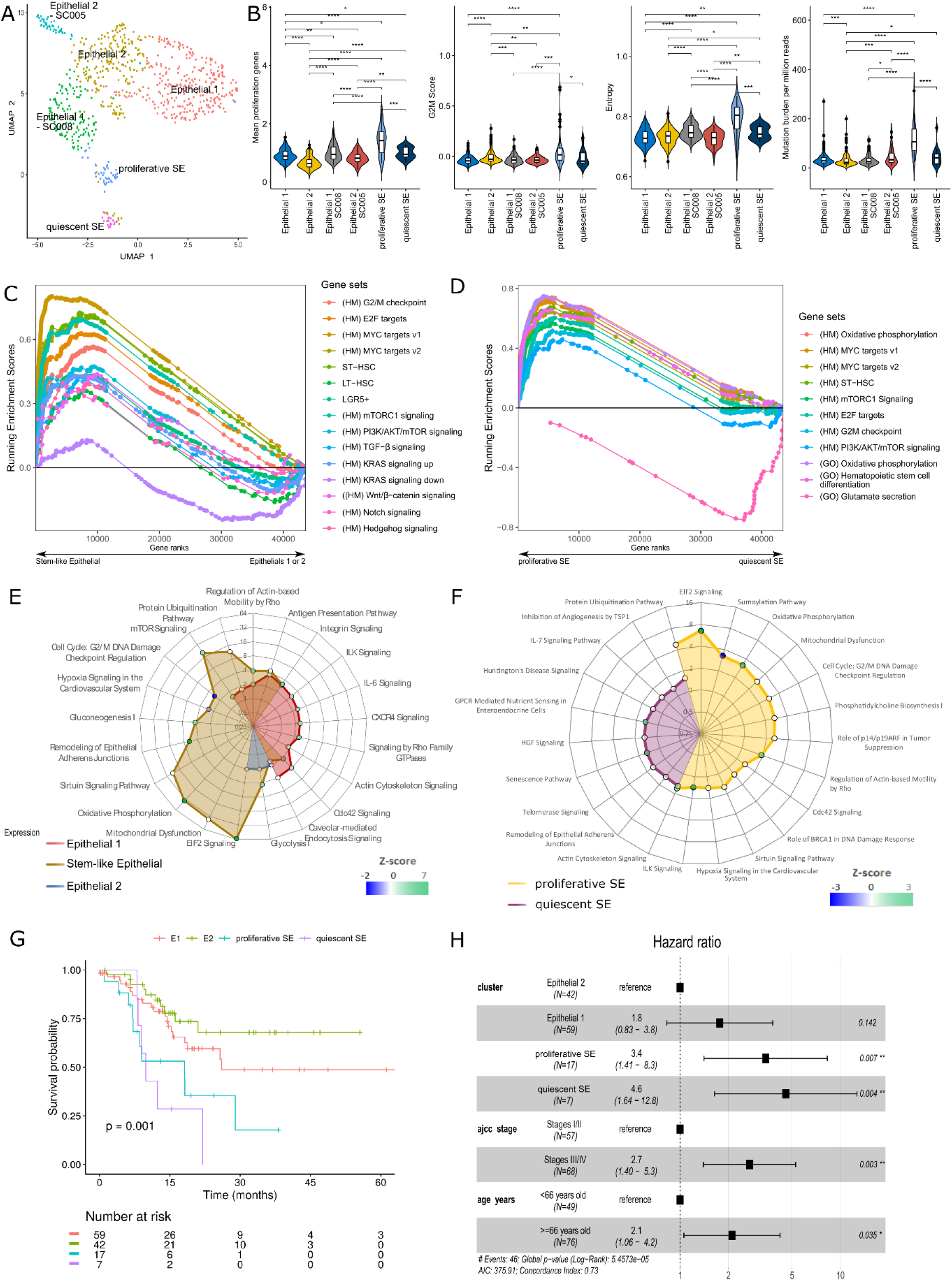
Two types of Stem-like Epithelial cells are identified. A) Six epithelial clusters were identified by our analysis: Epithelial 1 (E1), E2, E1-SC008, E2-SC005, proliferative Stem-like Epithelial, and quiescent Stem-like Epithelial. B) Comparisons of entropy score, mean expressions of proliferation genes, G2M cell cycle score, and tumor mutation burden between epithelial clusters. C-D) Gene Set Enrichment Analysis using gene sets from the MSigDB Hallmark v7.0 database. Statistical testing was performed by phenotype permutation test with FDR < 25% for each comparison. E-F) Differentially regulated pathways found with IPA based on the DEGs of E1, E2, and SE cell clusters, and differentially regulated pathways based on DEGs from the comparison between pSE and qSE. G) TCGA gastric cancer cohort patients were classified based on their similarity to the epithelial clusters to evaluate the association between epithelial cell type and overall survival. The number of patients that were E1, E2, pSE, and qSE -like were 101, 76, 24, and 9, respectively. The p-values from the pairwise log-rank tests were Bonferroni-corrected. H) Multivariate analysis of overall survival using the Cox proportional hazards regression, accounting for cell type, age, and stage.

We next characterized the two new clusters for their proliferative capacities using the G2/M cell cycle scoring (G2M) with Seurat [34], and proliferation-related gene set (GO:0008283). We found the two sets to be distinct with the first having both a high G2M score and high mean proliferation gene expression, indicating high proliferative activities (Figure 2B). Thus, we provisionally labelled the first as proliferative SE (pSE) and the second as quiescent SE (qSE). We then calculated the differentiation potency (signaling entropy) and pSE cells again scored the highest entropy score, followed by qSE, which supported their stem-like nature. We also quantified the mutation burden and found pSE to harbor the most point mutations (Figure 2B). It was the only epithelial cell cluster to have duplicated or deleted chromosomal regions (Figure S2E-H).

We next employed GSEA [41] to discover enriched gene sets. When comparing the stem-like clusters (pSE and qSE) to other epithelial cells (E1 and E2), the stem-like clusters revealed enrichment in stem-cell-related pathways such as KRAS signaling, Wnt/β−catenin signaling, Notch signaling, and Hedgehog signaling (Figure 2C). Proliferation related gene set targets, such as G2M checkpoint, E2F, MYC, short term hematopoietic stem cells (ST-HSC), long term hematopoietic stem cells (LT-HSC), and LGR5-positive stem cell signature genes, were also enriched. Using Ingenuity Pathway Analysis (IPA, Qiagen, GmbH), we found the stem-like cells to possess upregulated pathways such as EIF2 signaling, oxidative phosphorylation, and mTOR signaling, suggesting elevated metabolism and proliferation (Figure 2E). Comparing pSE and qSE, pSE was associated with higher levels of proliferation (mTORC1 Signaling, E2F targets, PI3K/AKT/mTOR signaling, oxidative phosphorylation) (Figure 2D). Similar results were obtained with IPA for the pSE cells. Meanwhile, qSE cells showed other differentially regulated pathways, including senescence that suggested them to be less proliferative (Figure 2F). The full list of DEGs and enriched pathways can be found in Tables S5-9.

We analyzed the correlation of the SE cells with patient survival as stem-related gene upregulation is associated with poor cancer patient survival [42]. We classified the TCGA GC patients [43] by their transcriptomic similarity to the different epithelial cell clusters, and then employed their survival status for survival analysis (Figures 2G,H). Univariate survival analysis showed that patients with high pSE or qSE signatures had poorer survival than patients that were E1 or E2 -like in pairwise comparisons (p < 0.05), except for E1 versus others (Figure S3). Comparing pSE and qSE, qSE-like patients had poorer survival than pSE-like patients. Similarly, multi-variate survival analysis adjusted for cancer stage and patient age showed that the pSE- and qSE-like patients had poorer survival than E2-like patients (Figure 2H). These results affirmed the negative impact of cancer stem-like cells on patient survival.

### pSE and qSE cells resembled IQGAP3^+^ isthmus stem cells and enteroendocrine cells respectively

We then investigated how our SE cells resembled the gastric stem cell populations previously reported. We found that many reported gastric stem cell markers MIST1 [24], LRIG1 [44], TROY [22], LGR5 [16], and AQP5 [27] were minimally expressed in all of our epithelial cell clusters (Figure S4A). However, STMN1 was upregulated in both pSE and qSE cells, as well as IQGAP3 and MKI67 in pSE (Figure 3A). These three genes have been reported [25,26] to mark actively cycling stem cells in the isthmus region of corpus gastric pits. Using the DEGs of reported gastric stem populations, we performed GSEA to investigate the resemblance of pSE and qSE to them. The upregulated gene sets in pSE strongly resembled the IQGAP3^+^ and MKI67^+^ stem cells, but qSE did not resemble any known gastric stem cells populations tested (Figure 3B).

**Figure 3:**
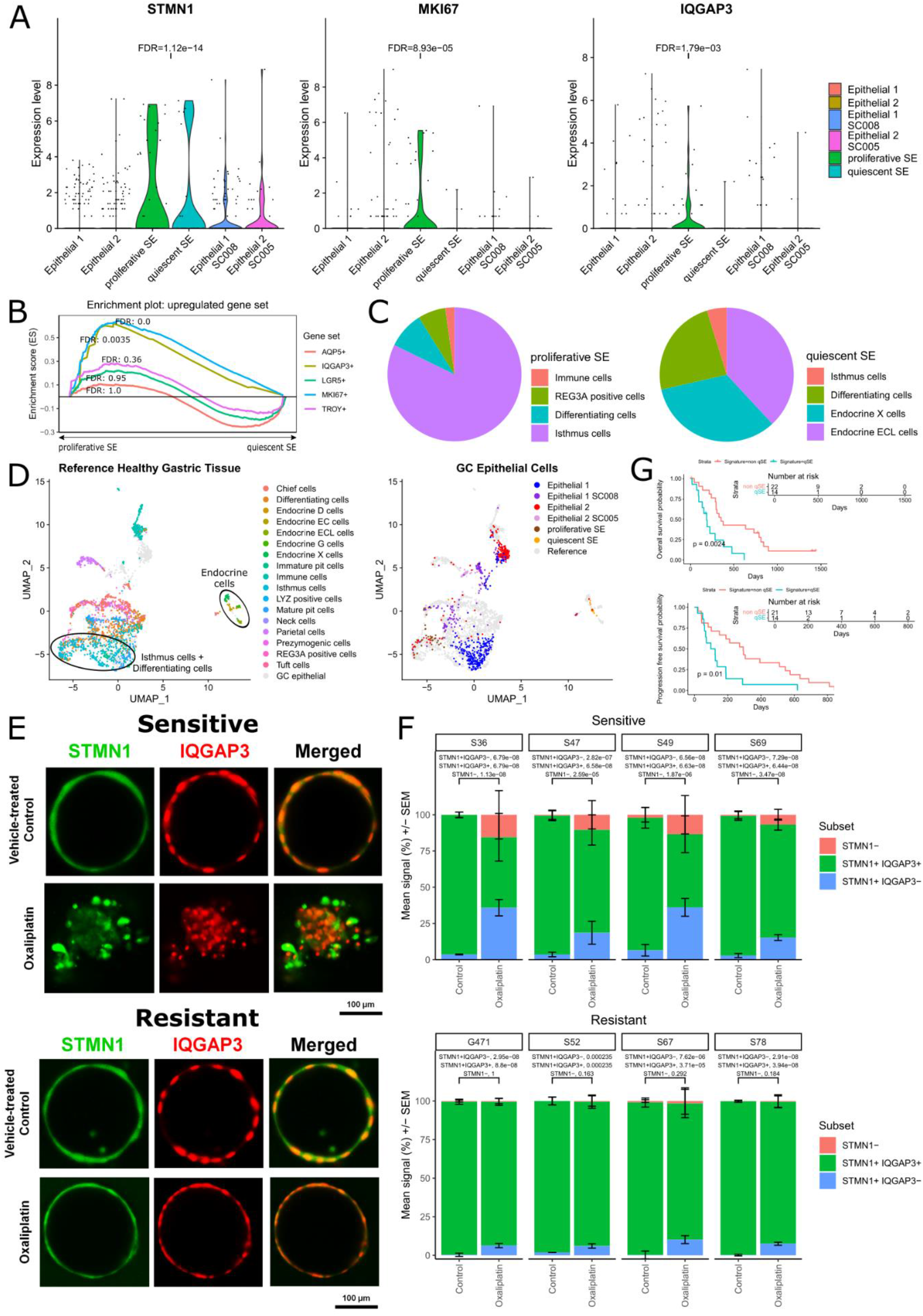
Proliferative SEs are similar IQGAP3^+^ MKI67^+^ gastric isthmus cells and quiescent SEs are similar to gastric enteroendocrine cells. A) Gastric stem cell markers (STMN1, IQGAP3, MKI67) expression in the different epithelial cell clusters. B) Comparison of pSE vs qSE cells in the enrichment of DEGs of previously identified gastric stem cell populations. C) Cell type prediction results by mapping pSE and qSE cells onto the healthy gastric cells by Busslinger *et al*. D) UMAP of data integrated pSE and qSE cells with healthy gastric tissue (Busslinger *et al*.) as reference. E) Immunofluorescence for STMN1 and IQGAP3 in patient-derived organoids before and after oxaliplatin treatment (1μM, 72 hours). F) Fractions of STMN1^-^, STMN1^+^IQGAP3^+^ and STMN1^+^IQGAP3^-^ cells in patient-derived organoids before and after oxaliplatin treatment. G) Progression free survival and overall survival analysis of GC qSE-like patients who underwent oxaliplatin treatment.

We also employed Seurat’s query projection functionality to map pSE and qSE onto the healthy gastric tissue map published by Busslinger *et al*. [45] to investigate their possible healthy origins. Most pSE cells were mapped to isthmus cells, supporting pSE’s isthmus origin. Intriguingly, qSE cells mapped to different enteroendocrine populations (Figure 3C and D). We further compared pSE and qSE to other published GC single cell data by integrating our data with Zhang *et al*.’s [32], the pSE cells overlapped with their metaplastic stem-like cell (MSC) and proliferative cell clusters (Figure S4B), and qSE overlapped with the enteroendocrine cluster. This suggested that qSE possibly originated from the enteroendocrine cells located in the gastric pit base. While the dedifferentiation of intestinal enteroendocrine cells to stem cells has been reported [46], similar occurrences in the stomach have not been reported.

### qSE-like STMN1^+^ IQGAP3^-^ cells are more resistant to oxaliplatin in chemosensitive GC organoids

As the pSE and qSE signatures were associated with poorer prognoses, we were interested in their response to common cancer treatments. Cancer stem cells are conventionally described as harboring dysregulated self-renewal pathways while remaining largely quiescent, with the latter property being a factor for chemoresistance [2]. pSE cells were characterized as highly proliferative and thus should be sensitive to chemotherapy, while qSE cells were quiescent and thus more resistant. To validate our hypothesis, we cultured GC derived organoids to test the survivability of different GC stem cell populations against chemotherapy.

The cultivated organoids were derived from separately acquired GC samples and tested for their drug resistance to oxaliplatin, a common platinum-based cancer medication that blocks DNA duplication (data not shown). Organoids selected from four sensitive and four resistant samples were stained for STMN1 and IQGAP3 (Figure 3E), and counted for three different populations (STMN1^+^ IQGAP3^-^, STMN1^+^ IQGAP3^+^, and STMN1^−^) before and after oxaliplatin treatment (Figure 3F). Prior to treatment, >90% of the cells were STMN1^+^ IQGAP3^+^, which is likely due to their high proliferative capacity. At 72 hours post treatment, the oxaliplatin-resistant organoids retained their structural integrity with minimal changes in cell population proportions. For the sensitive organoids, they lose structural integrity with a proportional decrease in STMN1^+^ IQGAP3^+^ cells while STMN1^+^ IQGAP3^−^ cells greatly increased in proportion. The survival of STMN1^+^ IQGAP3^−^ cells offers a plausible explanation for patients responding to chemotherapy but suffering from relapse due to STMN1^+^ IQGAP3^−^ cells surviving and contributing to the relapse. We further computed a qSE gene signature (Table S10) to segregate a local patient cohort who underwent oxaliplatin treatment and analyzed their survival prospects. Here we found patients similar to the qSE cells have both poorer progression free survival and overall survival (Figure 3G). Together, these results highlighted the chemoresistance of STMN1^+^ IQGAP3^−^ cells.

### Signals from epithelial stem-like cells contribute to immune evasion

Through tumor-immune cell interaction, tumor cells avoid immune eradication and influence patient prognosis [47]. Here we used CellPhoneDB [48] to predict the possible interactions among immune and epithelial cell clusters found in our samples (Figure 4A). We found more cross talks between immune cells and stem-like cells than non-stem like cells, and interactions are higher in tumor tissue than normal-adjacent. Of greatest interest are the immune suppressing and pro-tumor interactions, as they can contribute to the associated poor prognosis. Here we see that the pSE cells engaged in more immune suppressing interactions in both normal-adjacent and tumor tissues. More pro-tumor interactions were observed with the qSE, particularly at the tumor sites. These observations help explain the poorer survival of patients with the pSE and qSE signatures.

**Figure 4:**
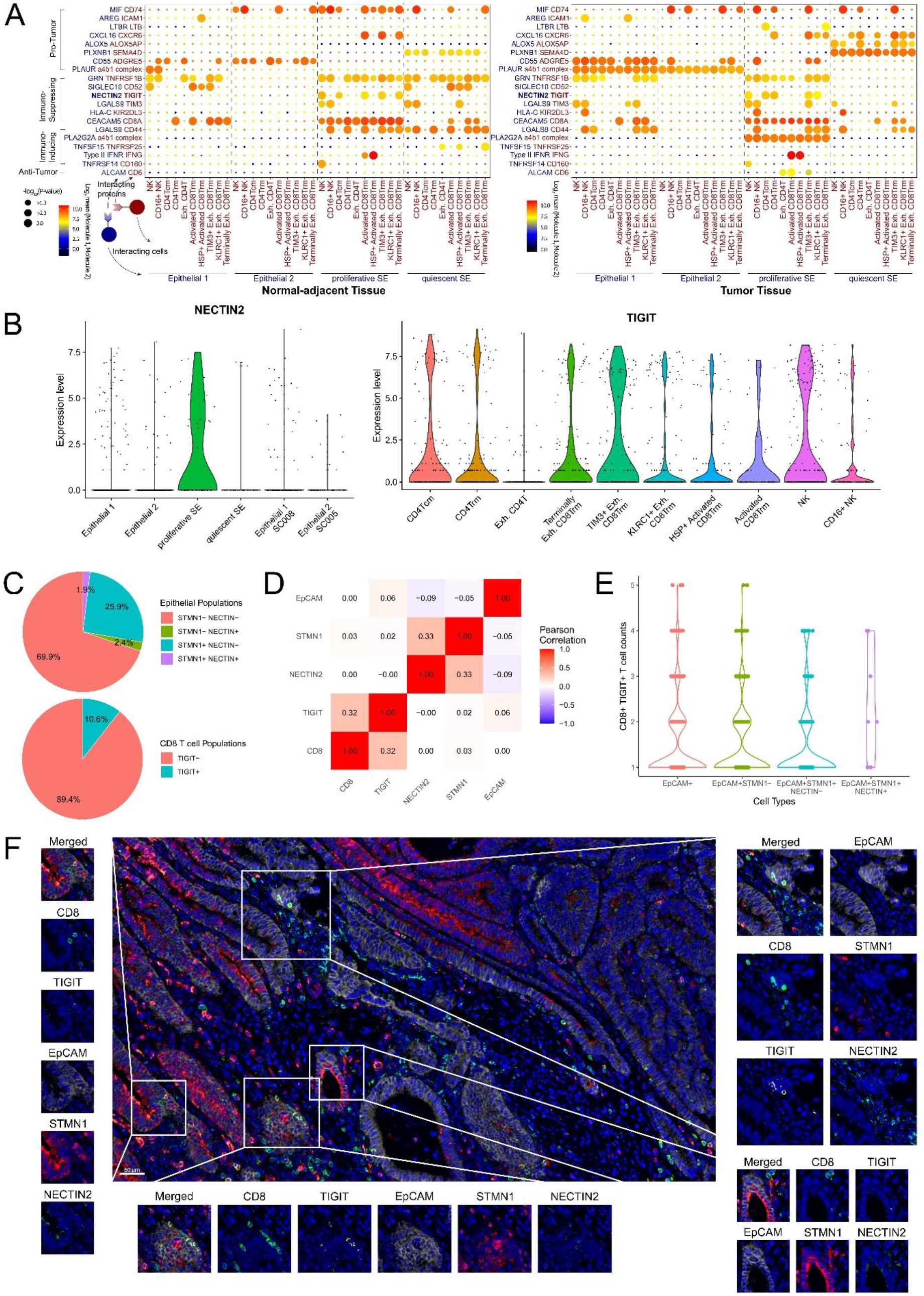
pSE and qSE inhibit CD8T and NK functions by expressing immune-suppressing ligands. A) Dot plots showing selected CellPhoneDB predicted ligand-receptor interactions between epithelial cell clusters and immune subsets. Shade represents mean expression of ligand-receptor pairs. Blue: expressed by epithelial cells; Red: expressed by immune cells. Negative log10 *p*-values are indicated by circle size. B) NECTIN2 and TIGIT gene expression in the epithelial clusters and immune cell clusters. C) Percentage breakdown of epithelial cell populations in IHC stained tissue slide by selected markers. D) Cell based correlation of detected IHC staining makers. E) Count distributions of CD8^+^ TIGIT^+^ T cells within 50 μm of an epithelial cell, grouped by different sub types. F) IHC staining slide image with regions of CD8^+^ TIGIT^+^ T cells and EpCAM^+^ STMN1^+^ NECTIN2^+^ epithelial cells in close proximity (50 μm) being highlighted.

Among the ligands engaged, many such as Galectin-9 (LGALS9)-TIM3 [49–51], GRN-TNFRSF1B [52,53], HLA-C-KIR2DL3 [54], and NECTIN2-TIGIT [55], can suppress T and NK cells in the tumor environment. Dysfunction in TIGIT-expressing CD8^+^ T cell is also linked to the expression of the NECTIN2 ligand by tumor cells [56]. In our data, NECTIN was specifically expressed in the pSE cluster among the epithelial cells (Figure 4B). We hypothesize that this interaction is a contributing factor to the poor survival of pSE-like patients.

We therefore employed multiplex immunohistochemistry staining on a GC tissue sample (SC007) to validate the predicted NECTIN-TIGIT interaction. The patient was chosen for the high proportion of pSE cells (Table S10). We stained the sample for EPCAM, STMN1, and NECTIN2 to locate the immune suppressive epithelial cells, and CD8 and TIGIT for the inhibited CD8^+^ T cells. To facilitate the analysis, we developed an automated computational workflow to locate and enumerate cells stained with different markers. Among epithelial cells stained by anti-EPCAM, STMN1^+^ NECTIN2^+^ cells only made up a small proportion (1.9%), consistent with their rare stem cell-like features (Figure 4C). The correlation between NECTIN2 and STMN1 was much higher (Pearson Correlation = 0.33) than their respective correlations to EPCAM, indicating that many STMN1^+^ stem-like epithelial cells expressed NECTIN. For the CD8^+^ T cells, 10.5% were TIGIT^+^ (Pearson correlation = 0.32, Figure 4D). Though NECTIN-TIGIT binding might be transient, we anticipate TIGIT^+^ CD8^+^ T cells to be in close proximity to NECTIN^+^ stem-like epithelial cells for effective suppression. We therefore used our algorithm to look for CD8^+^ TIGIT^+^ T cells located within 50 μm of EpCAM^+^ STMN1^+^ NECTIN2^+^ epithelial cells (Figure 4E and F). Indeed, we were able to find multiple of such pairs, showing that the TIGIT-NECTIN interaction could be an important pathway through which pSE cells achieved immune escape by inhibiting CD8^+^ T cell functions.

## Discussion

Though dietary changes and lower *Helicobacter pylori* infection rate have somewhat reduced its occurrence, GC continues to be one of the most common cancers. Furthermore, frequent relapses and chemo resistances complicate treatment and depress survival rates. Here, we used single cell RNA-seq to reveal a heterogeneous collection of epithelial cells in GC samples. We found two stem cell-like clusters, pSE and qSE, that were able to form oxaliplatin resistant organoids in tissue culture. The pSEs resembled gastric stem cells from the corpus isthmus region, sharing the same markers (STMN1^+^ MKI67^+^ IQGAP3^+^) and proliferation propensity as their healthy counterparts (Figure 5). They also highly expressed immunosuppressing molecules such as galectin-9 and NECTIN, which inhibit cytotoxic T and NK cells, which contribute to immune suppression in tumor. qSEs expressed STMN1 but not IQGAP3, appeared quiescent, and resembled gastric enteroendocrine cells which do not have known stem cell activity. We postulate that qSEs arose from dedifferentiated enteroendocrine cells after injury, a phenomenon previously described in small intestine [46]. Survival analysis with pSE and qSE expression signatures linked them to poorer survival than signatures of other GC epithelial cells.

**Figure 5:**
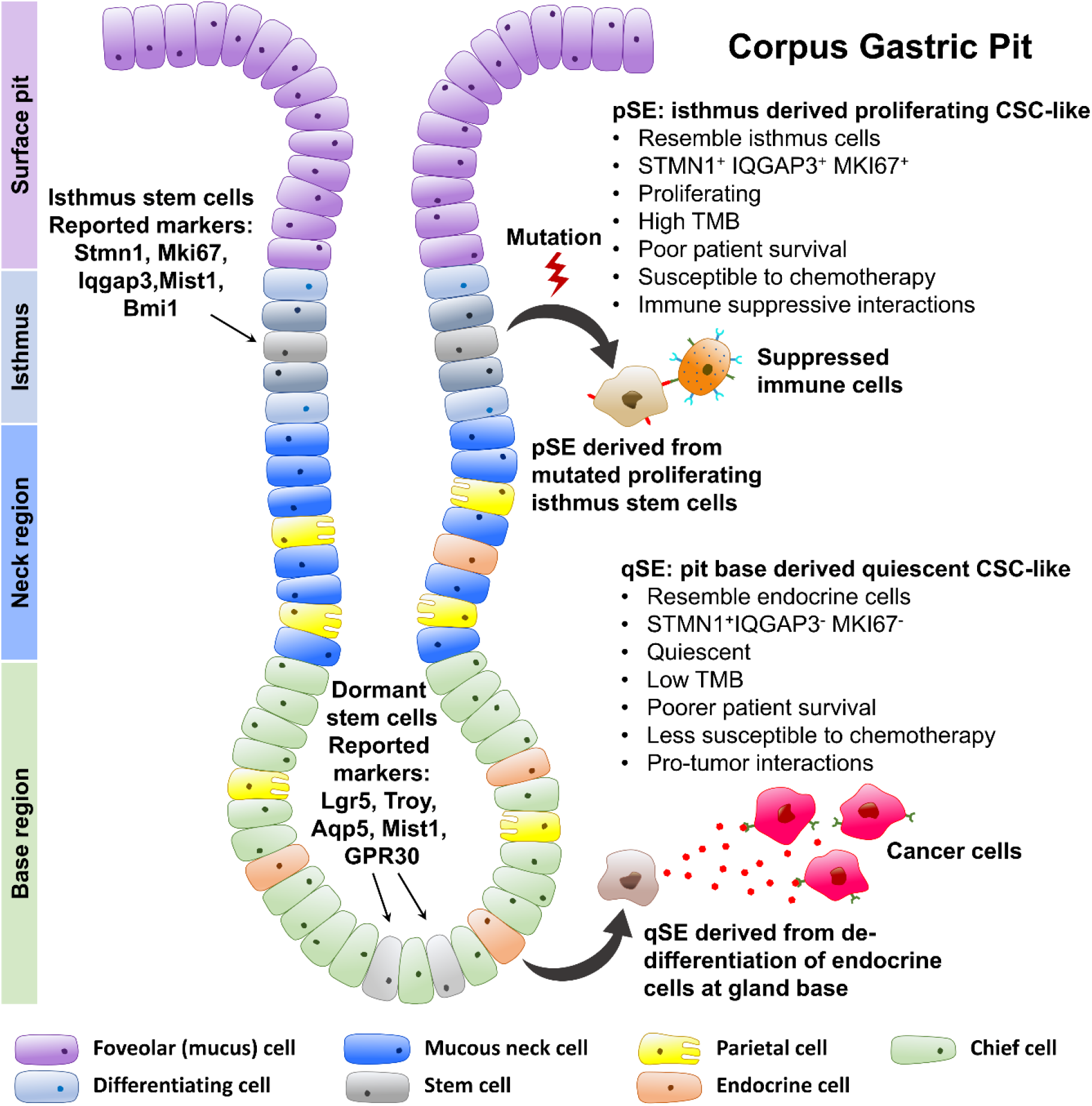
Cross section of a gastric corpus gland describing the different cell types lining the gland. The different reported healthy gastric stem cell populations and their markers are highlighted. From this study, proliferative SE cells appear to have derived from the proliferating stem cells in the isthmus region while quiescent SE cells resembled the enterendocrine cells found near the gland base.

The cultured organoids from GC samples showed that the pSE-like STMN1^+^ IQGAP3^+^ cells and the qSE-like STMN1^+^ IQGAP3^−^ cells can both be resistant to oxaliplatin, a chemotherapy that targets DNA replication. The large proportion by which the pSE-like STMN1^+^ IQGAP3^+^ cells made up the untreated organoids highlighted their proliferative nature. Active division increases the chance of spontaneous random mutations and consequently the risk of cells developing treatment resistance. Our results indeed showed that the STMN1^+^ IQGAP3^+^ cells possessed different levels of chemoresistance among patients, supporting our hypothesis that there are additional mutations accumulated within the pSE-like population. Once chemoresistance develops, the highly proliferative pSE-like population can contribute to drug-resistant relapse, which further worsens survival prospects. On the other hand, the qSE-like STMN1^+^IQGAP3^−^ cells are quiescent and therefore resistant to chemotherapies that target cell divisions. They are possibly reactivated later, leading to relapses. This can explain the observed poor progression free survival of qSE-like patients who underwent chemotherapy.

The pSE and qSE cells also expressed large numbers of immune-modulating molecules, of which many inhibit CD8^+^ T and NK function and possibly contribute to immune escape. We observed the expression of galectin-9 (LGALS9) in both pSE and qSE, and confirmed the potential immune-suppressive action of pSE-like STMN^+^ NECTIN2^+^ epithelial cells on TIGIT^+^ CD8 T cells by measuring their proximities in IHC staining. Our results suggest the potential benefits of immune checkpoint blockade such as TIM-3 [57] and TIGIT [58]. The identified pSE and qSE signatures can be used to screen for patients that can potentially benefit from these immunotherapies.

Therapy resistant CSCs are an ongoing concern in cancer treatment due to their role in relapse and poor survival. This makes it an imperative to target these cells to improve patient outcomes. Our results expectedly showed proliferating CSCs to be vulnerable to chemotherapy, but there were also samples that acquired chemoresistance. This development re-iterates the importance of targeting CSCs, as they are the proliferative source for cancer growth and mutations. Quiescent CSCs, by nature, are more chemoresistant and thus likely to survive conventional treatment. Therefore, the presence of different CSC populations necessitates additional elimination strategies. Moreover, the plasticity that enables tumor cells to dedifferentiate into CSCs poses additional challenges [59]. Strategies under investigation include the aforementioned immune blockade therapy, antibody-drug conjugate targeting CSC-specific markers, and targeting the stem cell niche by blocking signals such as WNT [6]. For quiescent CSCs, a proposed approach involves “waking” them up to increase their susceptibility to chemotherapy. A combination of these approaches is likely to be needed to increase overall efficacy, especially in conjunction with genotypic or phenotypic signatures that can inform on the nature of tumor cells present to guide therapy design.

## Author contribution

Jinmiao Chen conceptualized and supervised the study. Kok Siong Ang, Hong Kai Lee, Jingjing Ling, and Jinmiao Chen wrote the manuscript. Jimmy So and Wei Peng Yong recruited patients, collected samples, and provided critical insights. Vivien Koh optimized and performed the tissue dissociation process, generated the organoids from surgical specimens and maintained the cultures, and performed drug treatment experiments and whole mount immunofluorescence staining of the organoids. Calista Wong, Mai Chan Lau, Ze Ming Lim, Charles Antoine Dutertre, Anis Larbi, and Jinmiao Chen performed the FACS staining and index sorting. Michelle Goh, Alicia Tay, Josephine Lum, and Shanshan Wu ran single-cell smart-seq2. Jinmiao Chen, Marion Chevrier, Hong Kai Lee, Kok Siong Ang, Xiaomeng Zhang, Jingjing Ling, Jia Chi Tan, Nicole Lee, Carolyn Tang, and Hang Xu performed data analysis. Jia Chi Tan, Hong Kai Lee, Xiaomeng Zhang, Kok Siong Ang generated the figures. Sergio Erdal Irac, Chun Jye Lim, Sherlly Lim, and Joe Poh Sheng Yeong performed IHC staining. Samuel Chuah Wen Jin and Valerie Chew performed the FACS data gating. Michael Poidinger, Amit Singhal, Matthew Ng, Shanshan Wu, Patrick Tan, and Florent Ginhoux provided scientific advices.

## Acknowledgements

We would like to thank NMRC OFYIRG grant for funding this study; SIgN Immunomonitoring platform for running FACS and single-cell RNA-seq. SIgN Immunomonitoring platform is supported by a BMRC IAF 311006 grant and BMRC transition funds #H16/99/b0/011. We would also like to thank Fiona CHIA from the Microscopy Core Facility at the Cancer Science Institute of Singapore for her assistance with confocal microscopy.

## Supplemental methods

### Tissue dissociation and single cell sorting

All fresh tissues were first collected in cold FACS buffer (5% FBS and 2 mM EDTA in PBS) and then rinsed with cold HBSS (Sigma-Aldrich, MO). After mincing, the tissues were digested for 1-2 hours at 37 °C in 300 U/ml collagenase (Sigma-Aldrich) and 100 U/ml hyaluronidase (Sigma-Aldrich). Thereafter, the cell suspension was filtered and the cells were pelleted at 300 x g for 5 minutes at 4 °C. Cells were re-suspended and washed twice in cold FACS buffer. Cell viability was determined using trypan blue (Invitrogen, CA). Cells were again pelleted and subsequently re-suspended in a cold blocking buffer. An antibody cocktail was then added and incubated for 20 minutes on ice before FACS sorting. The stained cells were then sorted using the BD FACSAria III 3-Laser system. Stratified FACS-sorting was conducted to allow a 1:1 ratio of EpCAM^+^ and CD45^+^ cells collection; gating strategy is shown in Figure S5. The single cells were index-sorted and subjected to single cell RNA sequencing using the Smart-Seq2 protocol [1]. All samples were subjected to an indexed paired-end sequencing run of 2×151 cycles on an Illumina HiSeq 2000 and HiSeq 4000 system for about 1 million reads per sample.

### Single cell RNA raw data pre-processing

Reads from single-cell RNA-sequencing were mapped to the GRCh38 human reference genome GENCODE release 25 using STAR [2], and transcript per million reads (TPM) values were calculated by RSEM 1.3.0 [3].

### Seurat analysis and cell type identification

The TPM table was analyzed with the Seurat package [4]. Initial quality control was performed to remove cells with high mitochondrial activity (>50%), retaining 2,878 cells likely to be viable. SCTransform was performed on the gene expression values and the top 12,000 highly variable genes (HVGs) were selected for downstream analysis. Principal component analysis (PCA) was performed and the first 20 PCs were used to perform t-Distributed Stochastic Neighbor Embedding (tSNE) [5], Uniform Manifold Approximation and Projection (UMAP) [6], and clustering. Unsupervised Louvain clustering was performed at 0.8 resolution to cluster all the cells. To annotate the clusters, we computed the differentially expressed genes (DEGs) among the clusters using the LogNormalized values and the FindAllMarkers function with logfc.threshold=0 and bimodal statistical test. Pathways enriched with DEGs were identified using Ingenuity Pathway Analysis (IPA; Qiagen, GmbH) software. The same Seurat workflow was applied to the identified epithelial cells with the first nine PCs and sub-clustering at a resolution of 0.4.

### Datasets integration with Zhang et al

Seurat was used to integrate our SMART-Seq2 data with Zhang *et al*. [7]. Both datasets were log-normalized and 12000 HVGs were identified. We used the Seurat data integration protocol with the FindIntegrationAnchors and IntegrateData functions with default settings. The cells from Zhang *et al*. were annotated with gene markers published in the accompanying supplementary file.

### Cell of origin analysis

For cell of origin prediction, scHCL (version 0.1.1 https://github.com/ggjlab/scHCL/) was used with its default settings. The predicted results were processed manually to aggregate the cell types.

### Regulon analysis

Area Under the Curve (AUC) of regulon activities were calculated for epithelial cells using the SCENIC (Single Cell rEgulatory Network Inference and Clustering) [8] package with default parameters. Using Seurat, the AUCs of 624 regulons were directly used for PCA analysis, with the top 10 PCs used to generate a tSNE plot. Four clusters were identified at a resolution of 0.4. Differentially enriched regulons between clusters were identified using the Wilcoxon test.

### Gene Set Enrichment Analysis

Gene Set Enrichment Analysis (GSEA) [9] was performed using the Hallmarks v7.1 and C5–GO Biological Processes v7.1 gene sets available from the MSigDB database, long-term Hematopoietic Stem Cell (LT-HSC), short-term HSC (ST-HSC) gene sets [10], and the Lgr5-positive gene sets (GSE86603) [11]. Default settings of the GSEA software were used.

### Single-cell entropy analysis

The LandSCENT [12] was used to quantify differentiation entropy. The TPM values were log-normalized using the scater package. The DoIntegPPI function was used to integrate the normalized data with the human protein-protein interaction network (net13Jun12.m) defined by the package. Finally, the signaling entropy rate and estimated differentiation potency was computed with the CompSRana function.

### Top 500 proliferation and cell cycle genes

The expression levels of the human proliferation (GO:0008285, 1061 genes) and cell cycle (GO:0007049, 761 genes) genes were retrieved [13] and summed respectively across all cells for ranking. The top 500 highly expressed genes were then selected and used to compute the sum of expression for each cell.

### Epithelial-based classification of TCGA samples

To classify the TCGA GC samples by gene expression similarity with our epithelial cell clusters, the TCGA and our SMART-Seq2 (including all epithelial and non-epithelial cells) datasets were integrated using Seurat [14]. Both datasets were first normalized and scaled with the Seurat SCTransform normalization [15], and 3000 HVGs selected to identify the ‘anchors’ for datasets integration. After data integration, a global clustering of the combined data was performed to identify TCGA patients that clustered with the epithelial clusters. A total of 125 epithelial-like TCGA samples were selected to re-cluster with only the SMART-Seq2 Epithelial 1 (E1), E2, proliferative Stem-like Epithelial (pSE), and quiescent Stem-like Epithelial (qSE) cells for annotate by label transfer.

### Survival analysis of pSE and qSE -like TCGA patients

Overall survival curves were estimated by the Kaplan-Meier method and compared using the log-rank (Mantel-Cox) test. Patients who were still alive at last follow-up were censored. The Cox proportional hazards regression was used to relate risk factors to overall survival time. The cutp function from survMisc R package was used for the age risk factor to determine the optimal cut-off point.

### Tumor Mutation Burden

Non-synonymous variations of the coding genes were obtained using the GATK-3-4 RNASeq germline SNPs and indels workflow [16], with default options and reference. Variants found were annotated using snpEFF [17], with the same reference genome and annotation. The total genome coverage with minimum 4 pass-filter reads was surveyed using the GATK CallableLoci module. The tumor mutation burden (TMB) was derived by normalizing the total pass-filter non-synonymous variants that had minimal depth of 4 reads (counts) with the total genome coverage (per Mb) and total number of sequencing reads (per million reads).

### Immune cell clustering

Only cells identified as T and NK cells in the global clustering were re-clustered for analysis using Seurat. FACS data from patients SC016, SC017, and SC020 were used to identify CD4^+^ and CD8^+^ T cells among CD45^+^ cells and this information was used as the ground truth to annotate CD4^+^ T, CD8^+^ T, and NK cells. Clusters that lacked TCR expression and expressed GNLY were labelled as NK cells; clusters that overexpressed CD8A and CD8B were labelled as CD8^+^ T cells; the cluster that overexpressed CD4 and FOXP3 was labelled as Treg cells; all other clusters were labelled as CD4^+^ T. After confirming with the FACS data, one cluster was found to contain both CD4^+^ T and CD8^+^ T cells; this cluster was sub-clustered to separate them. Cells of each cell type were further re-clustered separately to further identify sub-clusters.

### Reconstruction of T cell receptor

T cell receptor sequences were reconstructed for all cells using the TraCeR [21] assemble function assemble function with “-s Hsap --loci A B G D” as input parameters. Clonal type and size were summarized on a per patient basis using the TraCeR summarize function with “-s Hsap --loci A B G D” parameters.

### Generation and maintenance of gastric cancer patient-derived organoids (PDO)

Human gastric cancer tissues were biopsied from each patient during surgical procedure and processed as described previously [22]. Briefly, tissues were first minced and then digested in phosphate-buffered saline containing 1 mg/ml collagenase (Sigma-Aldrich, Saint Louis, MI) and 2 mg/ml bovine serum albumin (BSA; Sigma-Aldrich) for 30 minutes at 37 °C. Cells were filtered, centrifuged at 300 × g for 5 minutes, resuspended in Matrigel (Corning Life Sciences, Corning, NY), and seeded into multiwell plates (Thermo Fisher Scientific). Cultures were maintained in gastric PDO culture medium [DMEM/F-12 supplemented with 10 mM HEPES (Thermo Fisher Scientific), 1× GlutaMAX (Thermo Fisher Scientific), 1× B27 (Thermo Fisher Scientific), 1× N2 (Thermo Fisher Scientific), 50 ng/ml epidermal growth factor (PeproTech, Cranbury, NJ), 100 ng/ml fibroblast growth factor (PeproTech), 1 nM gastrin (Sigma-Aldrich), 100 ng/ml noggin (PeproTech), 1 mM N-acetyl-L-cysteine (Sigma-Aldrich), 10 mM nicotinamide (Sigma-Aldrich), 10 μM Y-27632 (Sigma-Aldrich), 10% R-spondin conditioned medium, 50% Wnt3a conditioned medium] at 37 °C in 5% CO_2_ and passaged once every 7-10 days in 1:3 ratio.

### Drug treatment of PDO cultures

PDO cultures were treated with 10 μM oxaliplatin (Selleck Chemicals, Houston, TX) or dimethyl sulfoxide (DMSO; Sigma-Aldrich) as the vehicle control for 72 hours. After treatment, the cultures were processed for whole-mount immunofluorescence.

### Whole-mount immunofluorescence

Whole-mount immunofluorescence staining of PDO was performed as described previously [22]. Briefly, cultures were fixed with 3.7% formaldehyde (Sigma-Aldrich) for 15 minutes, permeabilised with 0.5% Triton X-100 (Sigma-Aldrich) for 20 minutes and then blocked with 2% goat serum (Thermo Fisher Scientific) for 1 hour. Samples were incubated with primary antibodies anti-STMN1 and anti-IQGAP3 (both from Novus Biologicals, Littleton, CO) at 4 °C overnight, washed with 0.01% Triton X-100, and then incubated with Alexa Fluor secondary antibodies (Thermo Fisher Scientific) at room temperature for 1 hour. Images were captured using Zeiss LSM 880 confocal microscope (Carl Zeiss AG, Oberkochen, Germany).

### Multiplex immunohistochemistry

Multiplex immunohistochemistry was performed on SC007 tumor slides using the Opal 7-colour kit (Akoya Biosciences, CA). The images were acquired using the Vectra 3.0 Automated Quantitative Pathology Imaging System (Perkin Elmer) with 4’,6-diamidino-2-phenylindole (DAPI) as the nuclear marker as previously described [24]. The antibodies used were anti-Human Stathmin (clone D1Y5A; Cell Signaling), CD8 (clone C8/144B; DAKO), TIGIT (clone 3106; Abcam), Nectin-2 (clone D8D3F; Cell Signaling), and EpCAM (clone VU-1D9; Invitrogen).

### Epithelial-based classification and survival analysis of qSE -like oxaliplatin treated patients

To classify the oxaliplatin treated patients by gene expression similarity, we first computed the qSE DEGs by comparing the qSE to the non-SE. We then employed CMAP to segregate the patients using the qSE DEGs as the gene signature. Statistically significant patients were retained for survival analysis. The survival curves were estimated by the Kaplan-Meier method and compared using the log-rank (Mantel-Cox) test. Patients who were still alive at last follow-up were censored.

### Cell-cell communication analysis

The CellPhoneDB2 [23] package was used to analyze ligand and receptor gene expression to predict cell-cell communications. The TPM table was analyzed using the cellphonedb method with default parameters, except that the minimum fraction of cells in a cluster expressing a gene was set to 0.3.

### Statistical analyses

The two-tailed Wilcoxon rank-sum test was used to compare nonparametric continuous variables, respectively. The Spearman’s Rank correlation was used to evaluate the strength and correlation (negative or positive) of a relationship between two non-parametric continuous variables.

### Cell specific ligand receptor quantification

We first used ImageJ to do cell segmentation on the DAPI stained image [25]. The raw image with blue fluorescence was transformed to grayscale (8-bit) and subjected auto thresholding to remove background noise. As we noticed that some regions had significantly lower fluorescence levels than the overall image, we employed Bernsen’s local thresholding method to ensure proper thresholding in the low fluorescence regions. We then applied the watershed algorithm to perform cell segmentation on the filtered image. The detected cells formed a bimodal distribution with the lower mode due to small segmented areas unlikely to be cells. Thus, we took the local minima as the cutoff to filter for actual segmented cells. Intensities of each marker for each segmented cell were calculated as marker value within cell divided by the cell’s area. We then used Rosin’s unimodal thresholding method to determine marker positive cells [26].

## Analysis of immune microenvironment

The immune compartment is one of the key components of the tumor microenvironment. Here we identified and characterized the different immune cell subtypes present in the samples. We first investigated the correlations between the percentage of immune cells and epithelial cells using the FACS output. We unbiasedly quantitated the live/dead CD45^+^ and CD3^+^ cells present in the tissues, except the first 5 patients recruited in this study, (i.e. Subjects SC001-SC005) where CD3^+^ information was not available. We found a significant positive correlation between the proportion of SE cells and percentage of CD45^+^ among the total live cells, within both normal-adjacent and tumor tissues (*p*<0.05; Figure S6A and B). This contrasts with the reported negative correlation between stem-ness and immune infiltration into solid tumors [1]. Indeed, we observed the antigen presentation pathway to be dysregulated in the SE cells (Figure 2E), which may have helped attract immune cells into the tumor. This suggests that despite a “hot” immune environment, the tumor cells were able to avoid immune eradication.

We next extracted the NK and T cells derived from the previous global clustering for re-clustering (Figure S6C) and subtype identification. Based on the expression of T cell receptor (*TCR*) genes, *CD8A*/*CD8B, GNLY, IL2RA* and *FOXP3* genes, we identified CD4^+^ T, CD8^+^ T, regulatory T cells (Treg), and NK cell clusters (Figures S6C and D; Table S12). The cell cluster that did not express TCR but expressed *GNLY* was labelled as NK cells, while the cluster that expressed TCR and *CD8A*/*CD8B* was labelled as CD8^+^ T cells, and the cluster that expressed *IL2RA* and *FOXP3* was labelled as Treg cells [2]. The remaining cluster was labelled as CD4^+^ T cells.

We then re-clustered the CD4^+^ T, CD8^+^ T and NK cell clusters separately for subsets. The NK cells were sub-clustered into CD16^+^ and CD16^−^ NKs based on CD16 expression (Figure S6E), with the CD16^+^ sub-cluster labeled as CD16^+^ NK cells. Upon activation, CD16^+^ NKs lose CD16 expression and become mature CD16^−^ NK cells [3,4]. The CD16^+^ NK cells upregulated *CCL4, KLRF1, CRTAM, EOMES*, and *SH2D1A*, while the CD16^−^ NK cells upregulated *IL32, TNFRSF18*, and *GZMA*.

Analysis of the CD4^+^ T cells identified five sub-clusters. We found one cluster of central memory T (Tcm) cells marked by the expression of *SELL* and *CCR7* [5], a resident memory (Trm) clusters marked by the expression of *CD69, ITGAE*, and *CD44* [6–9], and one cluster of exhausted CD4^+^ T cells with downregulated TCR expression (Figure S6F). The TCR downregulation has been reported in CD4^+^ T cells during TCR activation, and is thought to de-sensitize them to avoid prolonged and excessive inflammatory response [10–12]. These exhausted CD4^+^ T cells may have lost their function as CD4^+^ helper T cells in terms of antigen recognition and cytokine production. The CD4Trm cluster overexpressed *CCL5*, which is linked to the recruitment of CD4^+^ T into tumor site [14,15]. They also overexpressed various T cell activation markers, such as *ICOS, CCR6*, and *CXCR6*, indicating some level of CD4T activities in the tumor.

The CD8^+^ T cells all appeared to be resident memory T cells (Trm) as they expressed *CD69, CD44*, and *ITGAE*, but not homing markers such as *CCR7* or *LEF1* [5,16]. Sub-clustering of these cells revealed 5 subsets (Figure S6G), among which two cell subsets overexpressed IFN-γ, indicating that these cells were activated and functional. These two sets of activated Trm cells differed in their expression of heat shock proteins. One subset overexpressed protein chaperones such as *HSP70, HSP27, HSP47, DNAJ*s, as well as polyubiquitin precursors *UBB* and *UBC*. These cells might had been stressed during the sorting process and were therefore removed from the downstream analyses. We also found two subsets of exhausted cells that overexpressed *KLRC1* and *TIM3* respectively. The presence of these exhausted T cells is consistent with a number of previous reports [17–19]. To investigate whether exhausted CD8^+^ were more enriched in tumor site, we used IHC staining on patient’s tumor and normal adjacent samples and found more CD8^+^ *TIM3*^+^ T cells in the patient’s tumor tissue compared to normal tissue (Figure S7A). We also found another cluster of terminally exhausted CD8Trm, which showed marked down-regulation of CCL5 and cytotoxic agents such as granzymes, perforin and NKG7, indicating their lack of functionality.

Utilizing the RNA-seq reads, we reconstructed the T cell receptor sequences using TraCeR [20]. The assembled sequences were then analyzed to quantify clonal expansion among the CD4 and CD8 T cells (Figure S7B). Among the different CD8 subsets in tumor tissue, 44% - 89% of each subset are unexpanded singletons (Figure S7B). This is broadly similar to the proportion of singletons observed in other different cancer tissues [18]. In general, no statistical difference in proportions of cells with different degrees of clonal expansion was found in all CD4 and CD8 subsets, when comparing between normal-adjacent and tumor tissues.

---

**Supplemental Figure S1:**
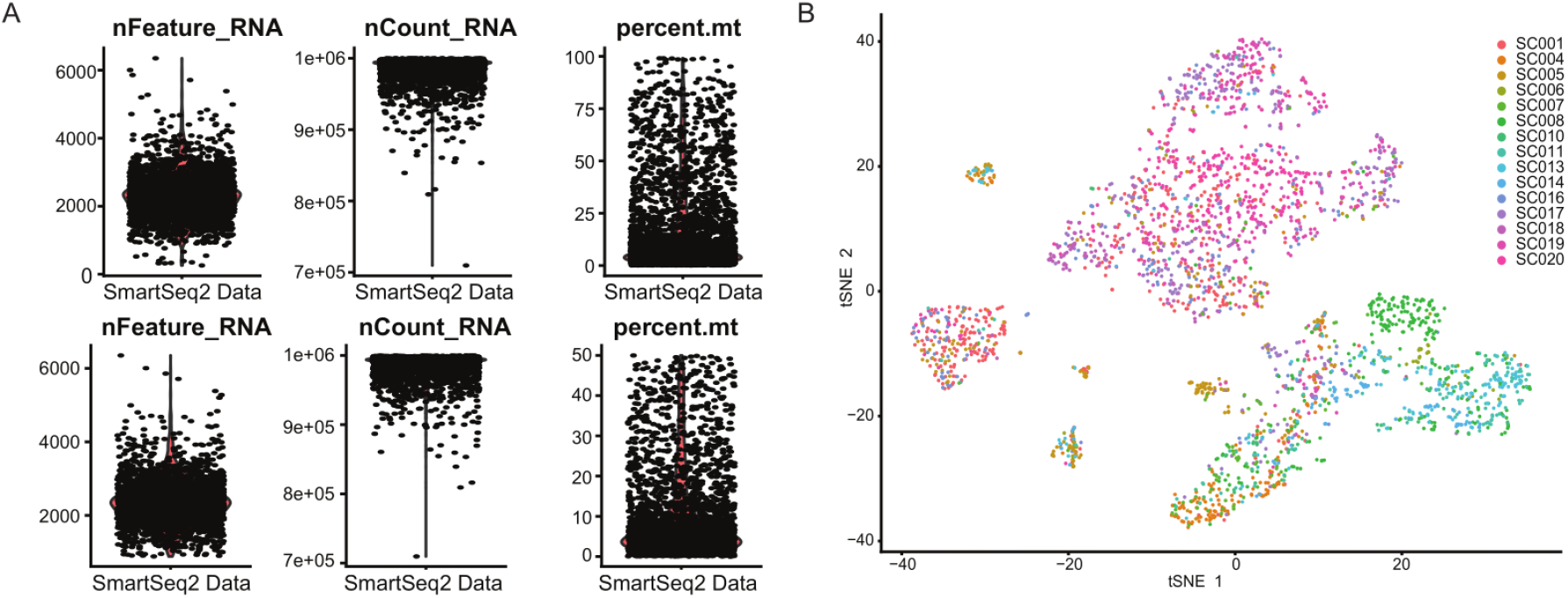
A) Violin plots showing cell quality before (left) and after (right) filtering sbased on three criteria, number of RNA feature per cell (nFeature_RNA) > 200 and < 5000 and percentage of mitochondria RNA count per cell (percent.mt) < 50%. B) tSNE plot of retained cells labeled by patient.

**Supplemental Figure S2:**
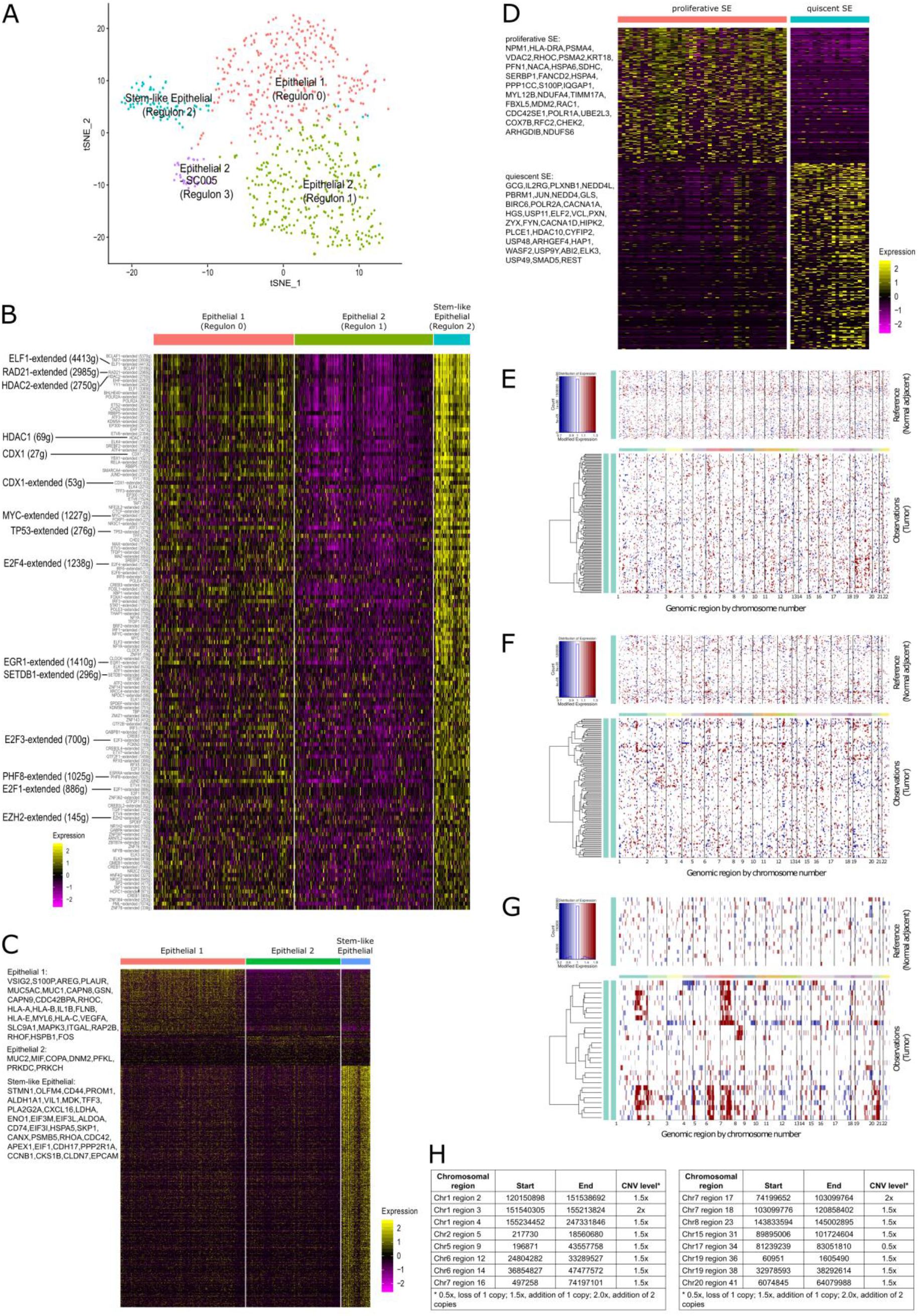
Identification of Stem-like Epithelials from Epithelial 1 and 2 clusters. A) Four epithelial clusters identified using regulon activity scores, including Regulon 0, 1, 2, and 3 clusters, each mostly corresponds to E1/E1-SC008, E2, SE, and E2-SC005, respectively. Each dot in the tSNE plot indicates an epithelial cell. B) Heatmap showing the differential regulon activities of Regulon 0 (E1/E1-SC008), 1 (E2), and 2 (ES) cells. C) Differential Expression Genes (DEGs) comparing E1, E2, and SE. D) DEGs comparing proliferative SE and quiescent SE. E-G) Predicted copy number variations (CNVs) by comparing (E) Epithelial 1 (E1), (F) E2, and (G) Stem-like Epithelial (SEA) obtained from the Normal-adjacent (Reference) and Tumor tissues. H) Level of the deletion or amplification event found in different chromosomal regions of Stem-like Epithelial A cells in tumor tissue (reference to normal tissue), filtered with a default posterior probability threshold of 0.5 that removed copy number variations (CNVs) that are not likely to be true CNV events.

**Supplemental Figure S3:**
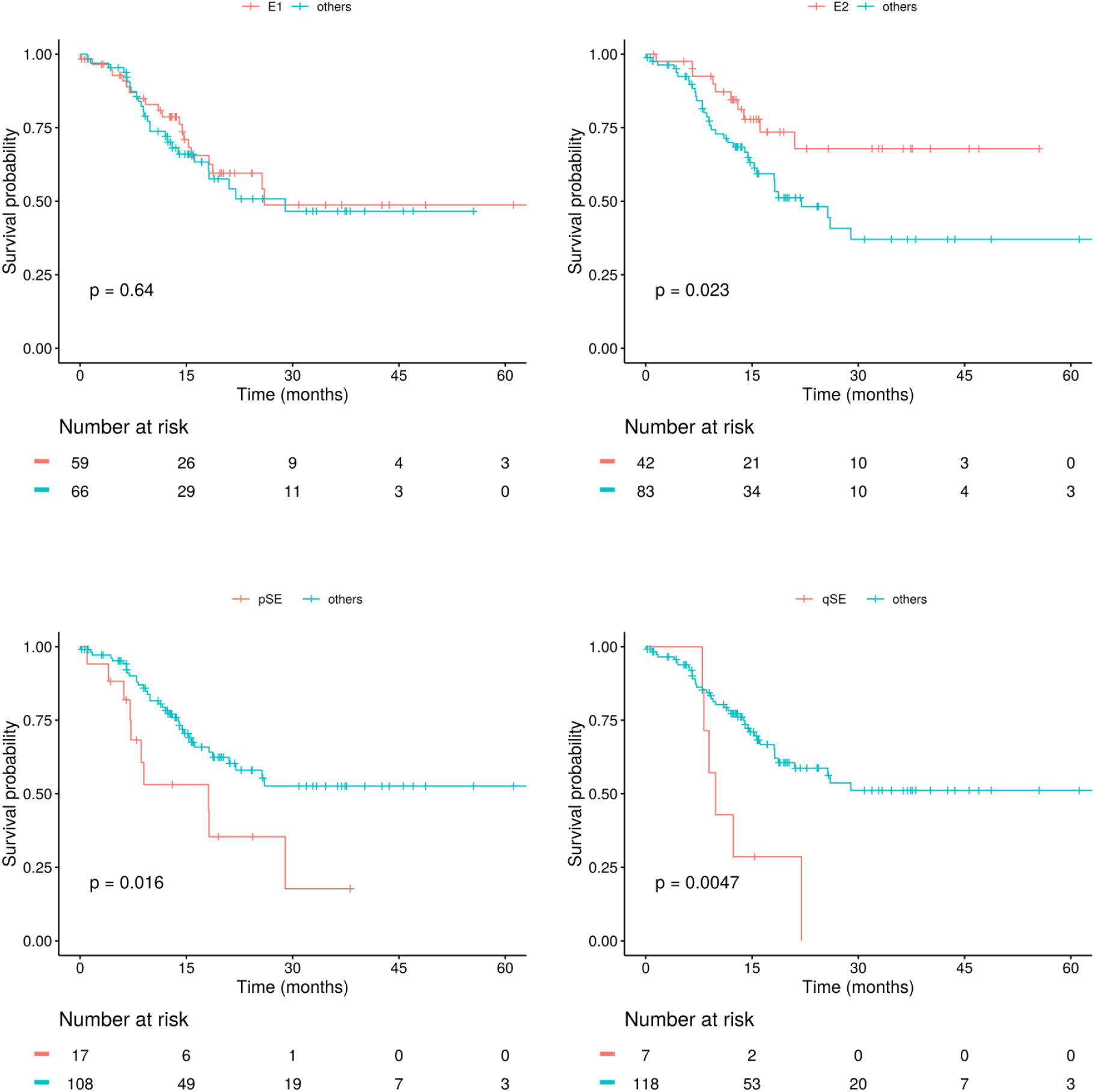
Kaplan Meier survival curves of TCGA patients classified as E1, E2, pSE, and qSE -like.

**Supplemental Figure S4:**
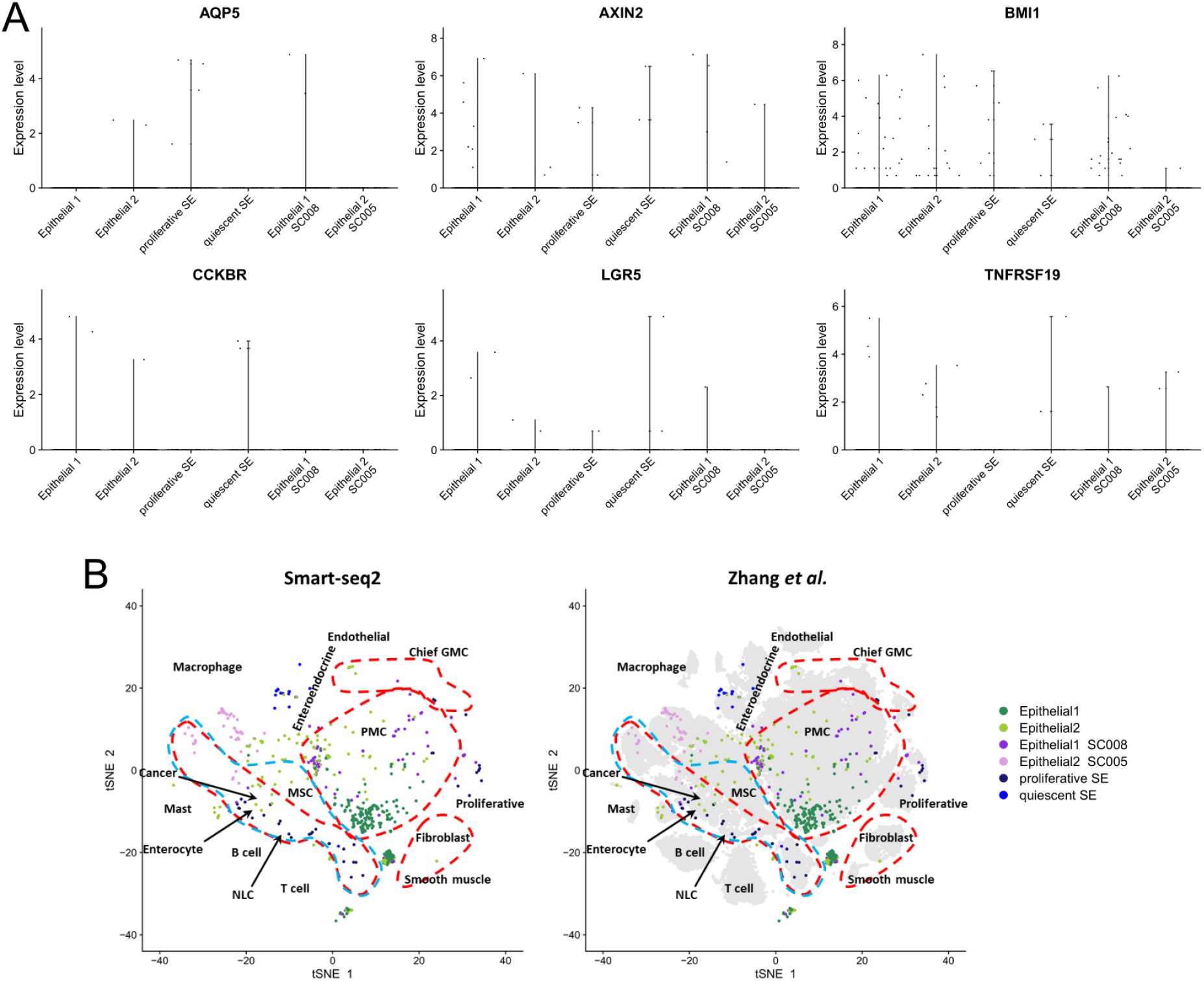
A) Gene expression level of other reported gastric stem cell markers. AQP5, AXIN2, BMI1, CCKBR, LGR5, TNFRSF19 (Troy). Mist1 (or Bhlha15) is not plotted due to it being filtered out during QC. B) Integration with dataset obtained from Zhang *et al*. and dashed lines represent areas of different cell types (green for MSC and red for epithelial and immune cells). GMC, antral basal gland mucous cell; MSC, metaplastic stem-like cell; NLC, neck-like cell; PMC, pit mucous cell.

**Supplemental Figure S5:**
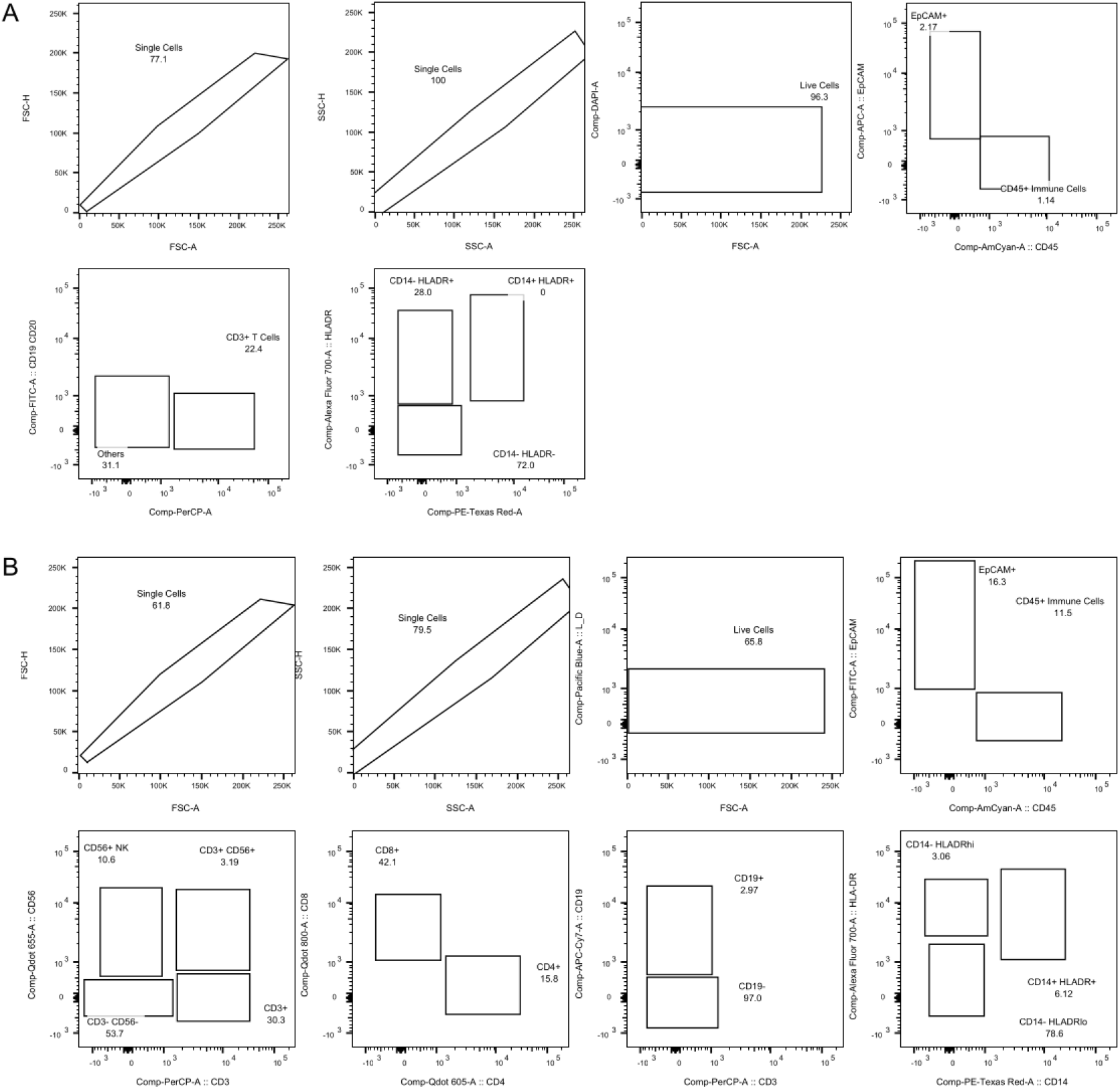
FACS gating strategies. (A) FACS Panel 2 and (B) FACS Panel 3 used in this study.

**Supplemental Figure S6:**
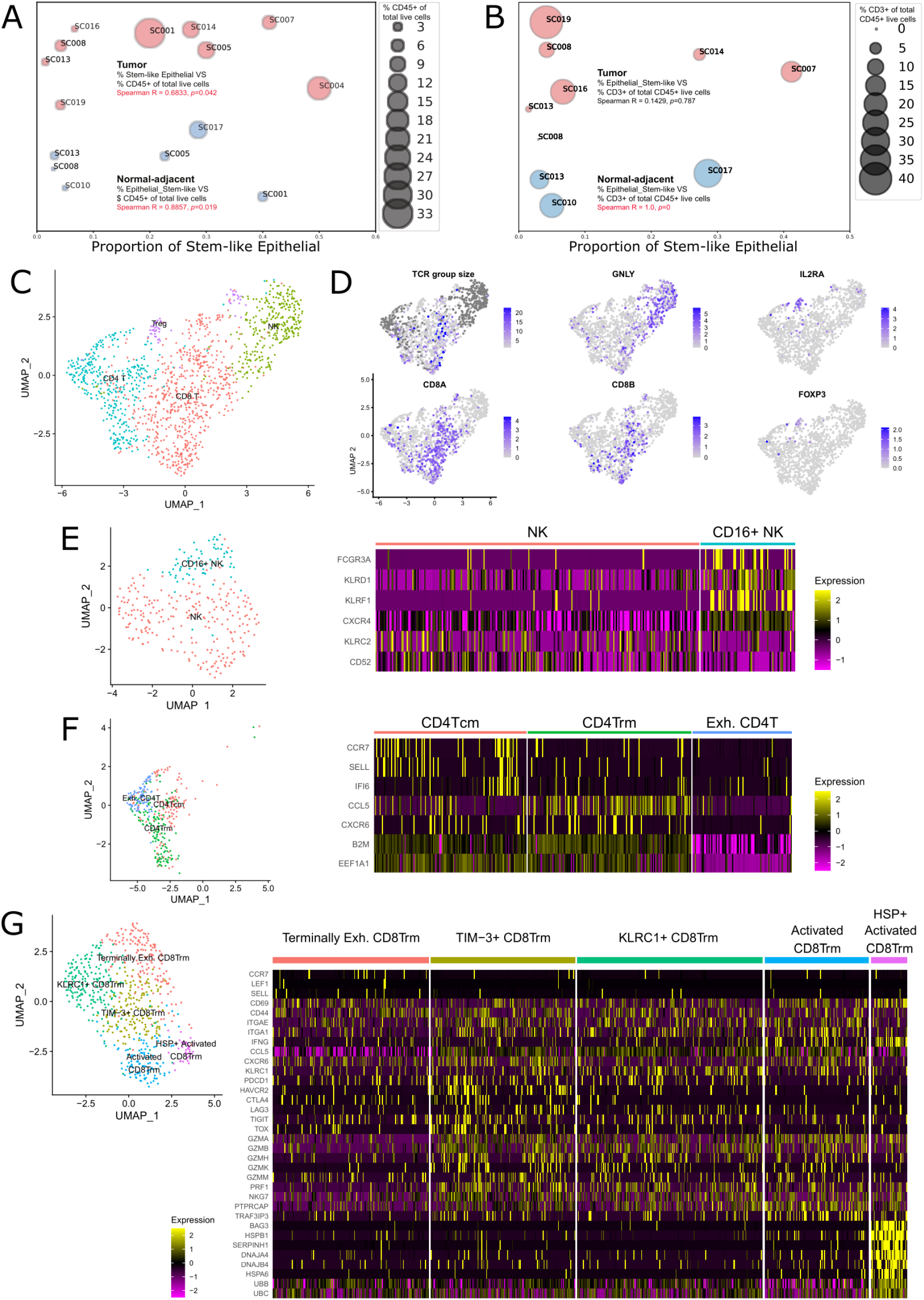
Dissection of immune heterogeneity and immune-signatures-based stratification for patient survival. A) Significant positive correlation between proportion of Stem-like Epithelial cell (of all the epithelial cells) and proportion of CD45^+^ immune cell (of all live cells) was found in gastric Normal-adjacent and Tumor tissues. B) Bubble plots shows significant positive correlation between proportion of Stem-like Epithelial cell (of all the epithelial cells) and proportion of CD3^+^ cell (of all CD45^+^ immune cells) was found in gastric normal-adjacent tissues (Blue bubbles), but not in tumor tissues (Red bubbles). C) Four major immune cell clusters identified, namely CD4^+^ T, CD8^+^ T, NK, and T-regulatory (Treg) cells. Each dot on the uMAP plot indicates a single cell. D) tSNE plots of all immune cells, colored by TCR group size, gene expressions of known markers for NK (GNLY), CD8^+^ T (CD8A and CD8B), regulatory T (IL2RA), and CD4^+^ T (FOXP3) cells. E) Two NK sub-clusters identified by known NK markers. F) Five CD4^+^ T sub-clusters identified by known CD4^+^ T markers. G) Four CD8^+^ Trm sub-clusters identified using gene expression values and the heatmap for known CD8^+^ T markers.

**Supplemental Figure S7:**
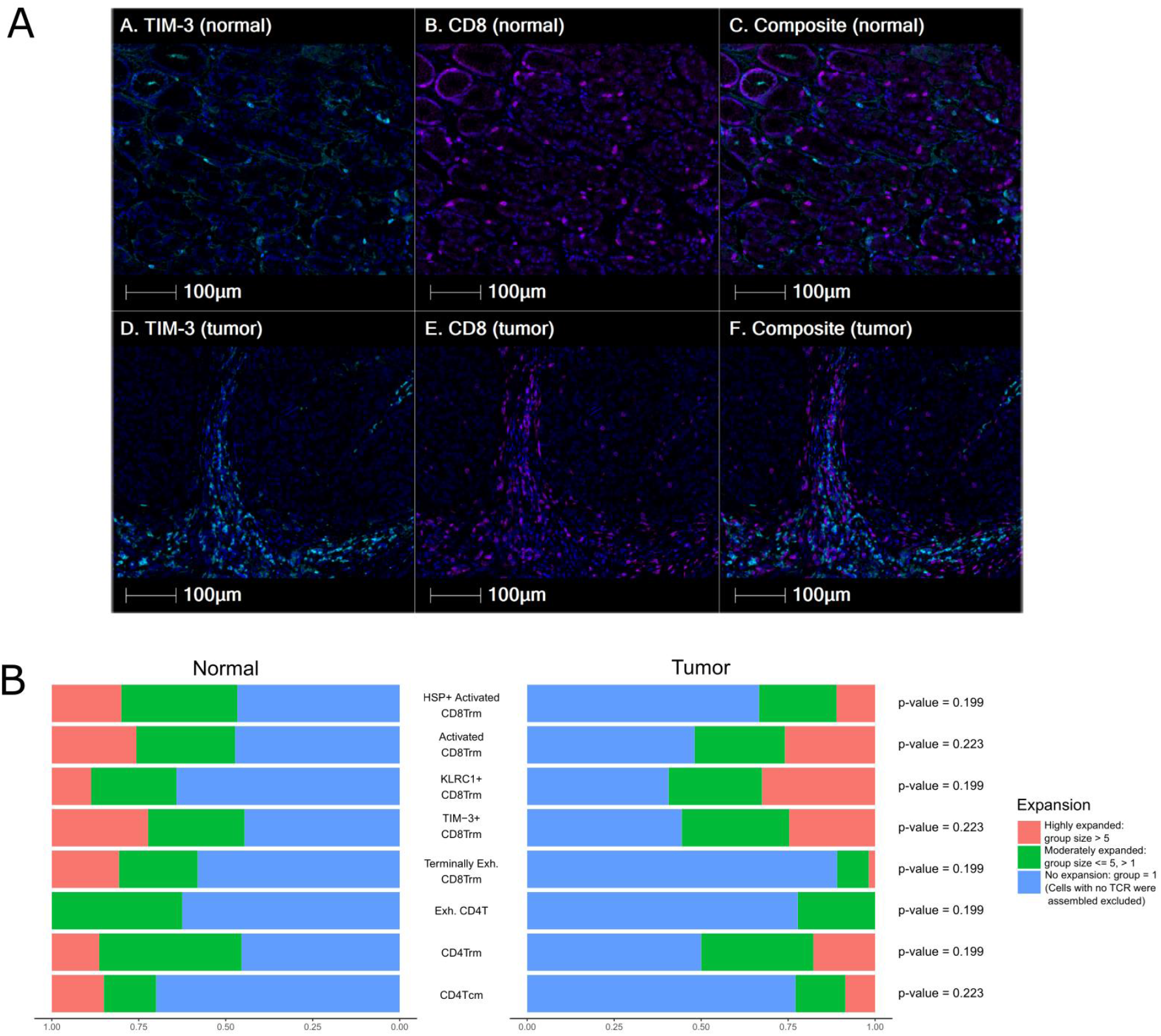
A) Immunohistochemistry staining of normal-adjacent and tumor tissues for an intestinal patient, SC019. TIM3^+^ and CD8^+^ cells were detected in both tissues. B) Distribution of T cell receptor clonal expansion patterns in CD4^+^ T and CD8^+^ T cell subsets. Normalized stacked bar plot for each subset show the fraction of cells with different degrees of clonal expansion, with respect to Normal-adjacent and Tumor tissues. Cells with unsuccessful TCR assembly were excluded.

## Notes

### Competing Interest Statement

The authors have declared no competing interest.

